# Single-cell profiling reveals tumor grade-dependent immune remodeling in *BRAF*^V600E^-driven glioneuronal tumors

**DOI:** 10.64898/2026.08.10.743180

**Authors:** Kittitach Sri-ngern-ngam, Philipp Müller, Anne Quatraccioni, Valentina Zschernack, Motaz Hamed, Rainer Surges, Susanne Schoch, Julika Pitsch, Albert J. Becker, Silvia Cases-Cunillera

**Affiliations:** Institute of Cellular Neurosciences II, Medical Faculty, University of Bonn, 53127, Bonn, Germany; Clinic and Polyclinic for Psychiatry and Psychotherapy, Medical Faculty, University of Bonn, 53127, Bonn, Germany; Institute of Neuropathology, Medical Faculty, University of Bonn, 53127, Bonn, Germany; Department of Neurosurgery, Medical Faculty, University of Bonn, 53127, Bonn, Germany; Department of Epileptology, Medical Faculty, University of Bonn, 53127, Bonn, Germany

**Keywords:** Glioneuronal tumors, microglia, T cells, Spp1

## Abstract

*BRAF*^V600E^ is the key driver variant in epilepsy-associated glioneuronal tumors (GNTs). These tumors often share MAPK/PI3K hyperactivation, a generally benign biological course and rare occurrence of malignant variants. We aimed to characterize the poorly defined immune cell milieu of GNTs with distinct biological behavior. We mapped cellular heterogeneity of the tumor microenvironment (TME) using single-cell transcriptomics on murine models of low-grade (LG-GNT; *BRAF*^V600E^/*AKT*^A^) and high-grade (HG-GNT; *BRAF*^V600E^/*AKT*^A^/*Trp53*^KO^) tumors, generated via intraventricular in utero electroporation (IUE). Furthermore, *ex vivo* functional assays with CSF1R-mediated myeloid depletion were utilized to assess the role of identified signaling molecules on tumor viability. We observed a striking, grade-dependent immunological dichotomy: LG-GNT exhibited a permissive niche with prominent surveillance by T cells and pro-inflammatory microglia. In contrast, HG-GNT TME was characterized by a restricted T cell infiltration, massively dominated by myeloid cell infiltrates. Differential gene expression analysis identified *Spp1* (osteopontin) as a key mediator of this immunosuppressive HG-GNT TME, exclusively expressed in microglia and border-associated macrophages (BAMs). Crucially, *ex vivo* functional assays demonstrated that recombinant SPP1 enhances tumor viability through a paracrine mechanism. These findings suggest fundamentally distinct immune activation (a) stimulated by aberrant MAPK/PI3K signaling in LG-GNT, versus (b) malignant tumor feature-driven, e.g. through necrosis in HG-GNT. In the latter, SPP1 signaling creates the immunosuppressive niche. Consequently, while modulating the pro-inflammatory niche may mitigate tumor-related epileptogenicity in LG-GNTs, targeting the SPP1-myeloid axis may restore anti-tumor immunity in HG-GNTs.

## Introduction

The immune cell architecture in epilepsy-associated neuroepithelial tumors so far remains largely enigmatic, despite the fact that these neoplastic lesions and their tumor microenvironment (TME) show prominent immune cell infiltrates. This holds particularly true for pleomorphic xanthoastrocytomas (PXAs) and gangliogliomas (GGs), both very frequent epilepsy-associated neoplasms [4, 20]. Clusters of neuronal cells have been described within the histopathological spectrum of PXA [18]. GGs generally show a benign neuropathological phenotype, but rare aggressive biological behavior has been reported [20, 22, 29, 36, 39]. In the 2021 WHO Classification of Tumors of the Central Nervous System, the term anaplastic ganglioglioma was removed owing to evidence that tumors previously diagnosed as such constitute a heterogeneous group of high-grade neuroepithelial neoplasms rather than a distinct entity [20]. Among these, PXA represents the most common diagnosis [33]. At the molecular level, the most frequent genetic mutation observed in GGs is *BRAF*^V600E^ [9, 16], a feature frequently shared by other epilepsy-associated tumors including PXAs (CNS WHO grade 2 or 3) [30, 38]. Furthermore, both GGs and PXAs often show PI3K-pathway activation [13, 32].

We have recently developed a neurodevelopmental brain tumor model in mice by intraventricular in-utero co-electroporation of clones of *BRAF*^V600E^ and a constitutively active (myristoylated) *AKT*, that recapitulates key neuropathological features of a low-grade glioneuronal brain tumor (GNT) resembling human GG [6]. By parallel inducing *Trp53*-loss, the resulting brain tumors acquired malignant features but retained the glioneuronal phenotype. Pathogenic mutations in *Trp53* have also been reported in a subset of high-grade PXAs, formerly called anaplastic PXAs [30]. Given these considerations, we refer to the tumors induced in mice for the present study as *BRAF*^V600E^-driven low-versus high grade GNTs (LG-GNT: *BRAF*^V600E^/*AKT*^A^; HG-GNT: *BRAF*^V600E^/*AKT*^A^/p53^KO^).

Here, we followed the hypothesis that the fundamentally different biological behavior of low-versus high grade GNTs is reflected by distinct immune cell-infiltrates in the tumors and the TME. To explore this, we applied single-cell transcriptomic profiling to map the heterogeneity of immune cell populations across these two murine models. By comparing LG-GNT and HG-GNT, we aimed to characterize grade-specific immune landscapes and identify potential microenvironmental mechanisms associated with epilepsy as well as tumor progression.

## Materials and methods

### Human samples

Histological images of human tumors were obtained from tissue resected from patients who underwent surgery at the Department of Neurosurgery of the University of Bonn.

### Mice

All animal procedures were planned and performed to minimize pain and suffering, and to reduce the number of animals. All experiments related to single-cell RNA sequencing were performed on B6.129P2-B6.129P2-*Trp53^tm1Brn^*/J mice backcrossed with CD1 mice for three generations to generate *Trp53^loxP/loxP^* mice on a CD1 genetic background. CD1 mice were obtained from Charles River Laboratories (strain code #022; RRID: IMSR_CRL:022), and B6.129P2-*Trp53^tm1Brn^*/J mice were purchased from The Jackson Laboratory (stock no. #008462; RRID: IMSR_JAX:008462). For all other experiments presented in this work, we used a CD1 genetic background. Mice were housed under a 12-hour light-dark cycle (7 AM - 7 PM), at 22°C ± 2°C and 55% ± 10% humidity, with ad libitum food, water, and nesting material (Nestlets, Ancare, Bellmore, NY), and ≥ one week of acclimation. Animals were individually housed post-surgery. All animals were maintained under sterile conditions on a 12-h light/dark cycle with ad libitum access to food and water. All animals were sacrificed between postnatal day 50-70 (P50-P70).

### Intraventricular *in utero* electroporation (IUE)

In utero electroporation (IUE) was carried out as previously described (Cases-Cunillera et al., 2022). In brief, pregnant female mice were operated on at embryonic day 14 (E14). Buprenorphine (0.05 mg/kg body weight) and ketoprofen (5 mg/kg body weight) were injected subcutaneously 30 minutes prior to the operation and anesthesia was induced by 3-4% of isoflurane and maintained at 1-2% throughout the procedure after the mice were deeply anesthetized. All DNA plasmids were prepared at a final concentration of 1.5 µg/µl and mixed with Fast Green FCF (Sigma-Aldrich) to allow visualization during injection. Following exposure of the uterus, the DNA solution was injected into the lateral ventricles of the embryos through the cerebral cortex using a glass capillary connected to a pressure-controlled microinjector (PICOSPRITZER^®^ III, Parker Hannifin). Electroporation was subsequently performed using the CUY21 SC Square Wave Electroporator (Nepa Gene) with 5 pulses of 30 V with a duration of 20 ms each. After injection and electroporation of the embryonic brains, the uterus was repositioned into the abdominal cavity, and the abdominal wall and skin were sutured. Postoperatively, mice were monitored daily and received subcutaneous ketoprofen injections for three consecutive days.

### Generation of plasmids

The DNA plasmids used for in utero electroporation were generated using PiggyBac (PB)–based vectors. Briefly, transgene sequences were inserted between the two PB terminal repeats of a donor PB plasmid (Wellcome Trust Sanger Institute; Cambridge, UK). pCMV-hyPBase constructs was acquired from the Wellcome Trust Sanger Institute. The *BRAF*^V600E^ transgene was generously provided by Dr. David Jones (German Cancer Research Center, Heidelberg). To generate the PB-CAG-mCherry-lynAkt construct (hereafter referred to as *AKT*^A^), the *Akt* coding sequence from myr-Akt1-pUSEamp (Addgene #17245; RRID: Addgene_17245) was PCR-amplified and subcloned into a plasmid containing the mCherry-T2A cassette (Addgene #72264; RRID: Addgene_72264). For single-cell RNA sequencing experiments, we used the murine ganglioglioma models previously described in [6]. For *in vitro* experiments, we generated novel ganglioglioma mouse models using in utero electroporation (IUE) in CD1 mouse embryos. CRISPR/Cas9-mediated knockout of *Trp53* was achieved by electroporation of the pX330-Trp53 plasmid (Addgene plasmid #59910), which encodes SpCas9 and a single-guide RNA targeting mouse *Trp53*.

### Single-cell preparation

Mice were decapitated, and brains were rapidly isolated and immediately immersed in ice-cold phosphate-buffered saline (PBS). Brain tissue was sliced into 300 µm sections, further microdissected, mechanically dissociated into small pieces, and transferred to ice-cold Hank’s Balanced Salt Solution (HBSS, Gibco). Single-cell suspensions were generated using the Neural Tissue Dissociation Kit (Miltenyi Biotec) according to the manufacturer’s instructions. Briefly, tissue pieces were transferred into C-Tubes containing Enzyme Mix X and Enzyme P and resuspended thoroughly. Subsequently, a second enzyme mix (Enzymes Y and A) was added. After gentle resuspension, samples were processed using the gentleMACS™ Octo Dissociator with Heaters (Miltenyi Biotec) using the preset Neural Tissue Dissociation program. Following dissociation, cell suspensions were filtered through 70 µm cell strainers and centrifuged. All further centrifugation steps were performed at 300 xg, 10 min, 4°C. The supernatant was discarded, and the resulting pellet was subjected to myelin removal using the Myelin Removal Kit (Miltenyi Biotec). After incubation for 15 min at 4 °C, samples were centrifuged, washed with PBS, and applied to LS columns (Miltenyi Biotec) according to the manufacturer’s protocol, retaining the myelin-positive fraction. The flow-through containing myelin-depleted cells was collected and washed with PBS. Subsequently, red blood cells were removed by incubation with Red Blood Cell Removal Reagent (Miltenyi Biotec) for 10 min at 4 °C. Cells were recovered by centrifugation and resuspended in HBSS (Gibco) for downstream applications.

### Single-cell RNA sequencing

Single-cell RNA sequencing (scRNA-seq) libraries were constructed using the 10x Genomics Chromium NextGEM Single Cell 3’ Reagent Kits v3.1, adhering strictly to the manufacturer’s protocols. To target a recovery of 1,500 nuclei per sample, nuclei were loaded onto a GEM Chip G. Sequencing was performed on an Illumina NovaSeq 6000 platform using a read configuration of 28/10/10/90 bp, yielding a mean sequencing depth of approximately 100,000 reads per nucleus.

### scRNA data processing

Raw scRNA sequencing data were processed in R using Seurat v4. Cells with fewer than 200 or more than 3,000 detected transcripts, or >10% mitochondrial reads, were excluded. After quality control, 9,496 cells were retained for downstream analysis. Data were normalized, scaled, and integrated using the standard Seurat workflow. Dimensionality reduction, clustering, and UMAP visualization were performed for cell-type identification based on canonical marker genes. Immune cell clusters were further subclustered and reanalyzed using the same workflow. Differentially expressed genes (DEGs) were identified using the Wilcoxon rank-sum test (min.pct = 0.1), followed by Gene Ontology enrichment analysis using PANTHER. Cell-cell communication analysis was performed using the CellChat R package [14] with CellChatDB.mouse as the reference database.

### Tumor cultures

For *ex vivo* culture experiments, tumor-bearing mice were sacrificed between P50-P70. Animals were deeply anesthetized with isoflurane and decapitated. Brains were sectioned and mCherry-positive regions were identified, microdissected and transferred into HBSS (Gibco #14170138). The isolated tissue was subsequently dissociated using the Neural Tissue Dissociation Kit (Miltenyi #130-092-628). According to the manufacturer’s protocol, tissue was enzymatically digested with enzymes P and A, and the resulting cell suspension was passed through a 70 µm cell strainer, followed by a 10-minute incubation with red blood cell lysis solution (Miltenyi #130-107-677). Cells were then washed, pelleted in PBS (Gibco #14190094; 300 xg, 5 min, 4 °C) and resuspended with Neurobasal medium (ThermoFisher #21103049) supplemented with 0.5x N-2 (ThermoFisher #17502048), 1x B-27 (ThermoFisher #17504001), and 0.5 mM GlutaMAX™ (ThermoFisher #35050061). Tumor cultures were plated and maintained at 37 °C in a humidified incubator with 5% CO₂.

### Mass spectrometry (MS)

Mass spectrometric analysis of condition medium (CM) from cell cultures was performed as previously described (Muller et al., 2024). Briefly, CM was collected, filtered, concentrated, and processed for proteomic analysis. Samples were prepared using the iST-NHS 96x kit (Preomics) and labeled with TMT10plex reagents (Thermo Fisher Scientific). LC-MS/MS analysis was performed using an Orbitrap Fusion Lumos mass spectrometer coupled to a Dionex Ultimate 3000 RSLC nanoHPLC system. Raw data were analyzed using Proteome Discoverer with Mascot searches against the Swiss-Prot mouse database. Gene Ontology enrichment analysis was performed using PANTHER [37].

### Histological examinations

Mouse tumor brains were collected between P50–P70, fixed in 10% paraformaldehyde (PFA), paraffin-embedded, and sectioned at 4 μm. Sections were stained with H&E for histological evaluation. For immunostaining, antigen retrieval was performed in citrate buffer (pH 6.0), followed by incubation with primary antibodies overnight at 4 °C. For immunofluorescence, fluorophore-conjugated secondary antibodies were used, and sections were mounted with Mowiol 4-88 (Roth #0718). For DAB-based immunohistochemistry, HRP-conjugated secondary antibodies and DAB substrate were used, followed by hematoxylin counterstaining.

### Western blot analysis

Tumor culture was treated with recombinant SPP1 (rSPP1) (ThermoFisher Scientific #120-35) and/or PLX3397 (Selleckchem #S7818) as indicated in the experiments. After 72 h of treatment, proteins were subsequently extracted by RIPA lysis buffer (50 mM Tris HCl pH 7.4, 150 mM NaCl, 5 mM EDTA, 1% nonidet P-40, 0.5% sodium deoxycholate supplemented with protease and phosphate inhibitors). Protein concentration was determined using a DC protein assay kit (Biorad #5000112EDU). Equal amounts of protein were separated by SDS-PAGE and transferred onto 0.45 μm PVDF membranes (Milipore #IPVH00010). The membrane was incubated with primary and secondary antibodies at the following concentration: rabbit anti-Iba1 antibody (1: 500, Wako #019-19741), mouse anti-β-actin (1: 8000, Abcam #ab6276), goat anti-rabbit IgG (H+L) conjugated to IRDye 800CW (1: 10000, LiCORbio #926-32211), and goat anti-mouse IgG (H+L) conjugated to IRDye 680RD (1: 10000, LiCORbio #926-68070). Signals were detected using the Odyssey CLx Imaging System and quantified with ImageJ. Protein expression levels were normalized to β-actin and expressed relative to the control condition.

### Cell viability assay

Cell viability was measured using the CellTiter-Blue^®^ Cell Viability Assay (Promega #G8080) according to the manufacturer’s protocol. In brief, tumor cells were prepared and plated in 96-well plates at a density of 1.0 × 10^4^ cell/ well and subjected to the indicated experimental treatments. At DIV4, CellTiter-Blue reagent was directly added to each well (1:10 dilution) and the plate was incubated at 37 °C for 2 h. Fluorescent signal was subsequently recorded using a microplate reader (excitation 560 nm., emission 590 nm.). Background fluorescence was subtracted and values were normalized to control samples.

### Statistics

All statistical analyses were performed using GraphPad Prism software using One-way ANOVA. Data are presented as mean ± SEM. Only statistically significant differences (p < 0.05) are shown in the figures.

## Results

### Dense immune cell infiltrates in GNTs

We aimed to systematically assess the GNT-associated immune landscape across GNTs with distinct biological behaviors. For this purpose, we studied low-grade (*BRAF*^V600E^/*AKT*^A^) and high-grade (*BRAF*^V600E^/*AKT*^A^/*Trp53*^KO^) mouse GNT models (hereafter referred to as LG-GNT and HG-GNT, respectively), developed via intraventricular in utero electroporation (IUE) as previously described [6]. As a control group (further referred to as Ctrl), we used mice subjected to IUE with a vector encoding a fluorescent marker. Hematoxylin and eosin (H&E) staining of Ctrl, LG-GNT, and HG-GNT tissues demonstrated that these models faithfully recapitulate the tissue architecture and histopathological features characteristic of human low- and high-grade GNTs (**Fig. 1A-B**). To evaluate major immune cell infiltrates within the TME, immunofluorescence staining for the pan-leukocyte marker CD45 was performed. While Ctrl brain tissue showed no detectable CD45^+^ signal, both LG-GNT and HG-GNT exhibited substantial CD45^+^ cell infiltration (**Fig. 1C**).

**Figure 1.**
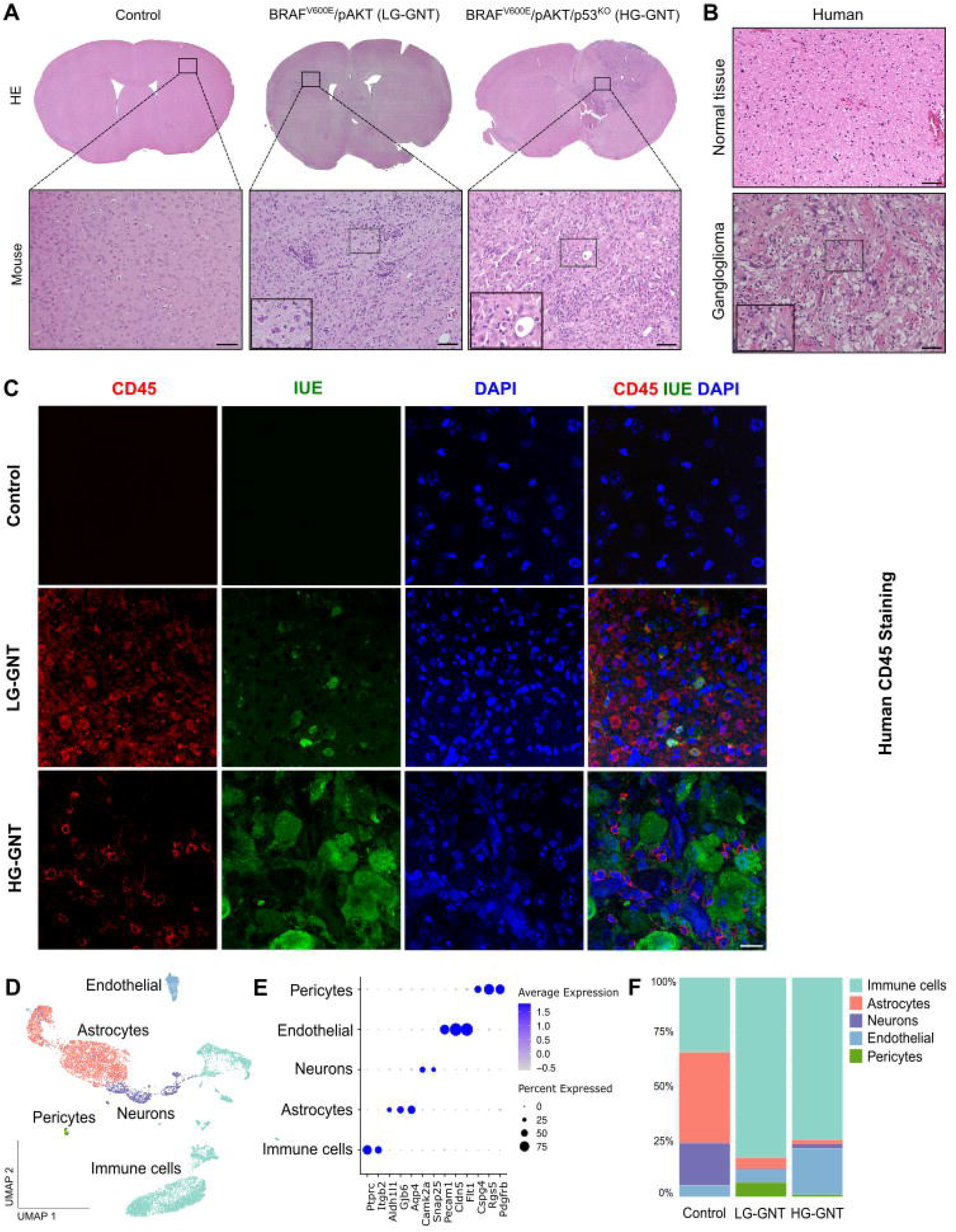
Histopathological and single-cell characterization of the tumor microenvironment in glioneuronal tumors (GNTs). **(A)** Representative Hematoxylin and Eosin (H&E) staining of IUE-brain cross-sections (top row) and high-magnification views (bottom row, scale bars = 100 μm) from Ctrl, low-grade glioneuronal tumor (LG-GNT; driven by *BRAF*^V600E^/*AKT*^A^), and high-grade glioneuronal tumor (HG-GNT; driven by *BRAF*^V600E^/*AKT*^A^/p53^KO^) mouse models. **(B)** Representative H&E staining of human tumor tissue sections, specifically GG representing LG-GNT, and PXA; representing HG-GNT (scale bars = 100 μm). **(C)** Immunofluorescence microscopy showing immune cell infiltration within the tumor area. Sections from Ctrl, LG-GNT, and HG-GNT tumor mice were stained against CD45 (red, pan-leukocyte marker), IUE (green, representing tumor cells introduced via In Utero Electroporation), and DAPI (blue, nuclei). Scale bar = 100 μm. **(D)** Uniform Manifold Approximation and Projection (UMAP) plot of integrated single-cell RNA sequencing (scRNA-seq) data, visualizing the distinct major cell populations within the microenvironment: immune cells, astrocytes, neurons, endothelial cells, and pericytes. **(E)** Dot plot displaying the expression profiles of canonical marker genes used for cell type annotation across the identified clusters. Dot size represents the percentage of cells expressing the specific gene, while color intensity indicates the average expression level. **(F)** Stacked bar graph quantifying the relative cellular composition (proportions of identified cell types) across the Ctrl, LG-GNT, and HG-GNT conditions. Notably, tumor-bearing models exhibit a marked expansion of the immune cell compartment.

We next employed single-cell RNA sequencing (scRNAseq) to comprehensively profile the cellular composition and transcriptional states of the immune cells in Ctrl (n = 3), LG-GNT (n = 3), and HG-GNT (n = 3) brain tissues. Unsupervised clustering of the integrated dataset identified five distinct clusters corresponding to neurons, astrocytes, endothelial cells, pericytes, and immune cells (**Fig. 1D**). These clusters were annotated based on the expression of canonical markers consistently validated across the literature; neurons (*Camk2a, Snap25*), astrocytes (*Aldh1l1, Gjb6, Aqp4*), endothelial cells (*Pecam1, Cldn5, Flt1*), pericytes (*Cspg4, Rgs5, Pdgfrb*) and immune cells (*Ptprc, Itgb2*, **Fig. 1E** and **Fig. S1**). In line with our CD45^+^ immunostaining data, our results showed a higher percentage of the total immune cell population within the TME in both GNT (LG-GNT: 82.07%; HG-GNT: 74.79%) models compared to Ctrl (33.10%) tissue (**Fig. 1F**). Moreover, the proportion of the immune cell population between LG-GNT and HG-GNT did not differ significantly (data not shown), indicating that GNTs are highly immunogenic regardless of the tumor’s biological behavior.

### Lymphocyte- to myeloid-dominant scRNA-seq signature drift in malignant GNTs

To comprehensively characterize the transcriptional heterogeneity of the immune microenvironment, we isolated and sub-clustered the immune cell population. UMAP-based dimensionality reduction identified eight distinct clusters spanning diverse myeloid and lymphoid cell lineages, which were annotated to specific cell types based on the expression of canonical marker genes (**Fig. 2A-B**, (Chen et al., 2018; Dean et al., 2024; Rajendran et al., 2023; Woolf et al., 2021; B. Zhang et al., 2024)). We identified resident microglia (*P2ry12*, *Tmem119*), alongside myeloid cells, including border-associated macrophages (BAMs; *Mrc1*, *Dab2*), infiltrating monocytes (*Ly6c1*), mast cells (*Ms4a2, Gata2*) and neutrophils (*S100a8*, *S100a9*) as well as a lymphoid cells consisting of CD3^+^ T cells (*Cd3e*), B cells (*Cd79a, Cd19, Ms4a1*), and natural killer cells (NK; *Nrc1, Klrb1c*, **Fig. 2B**).

**Figure 2.**
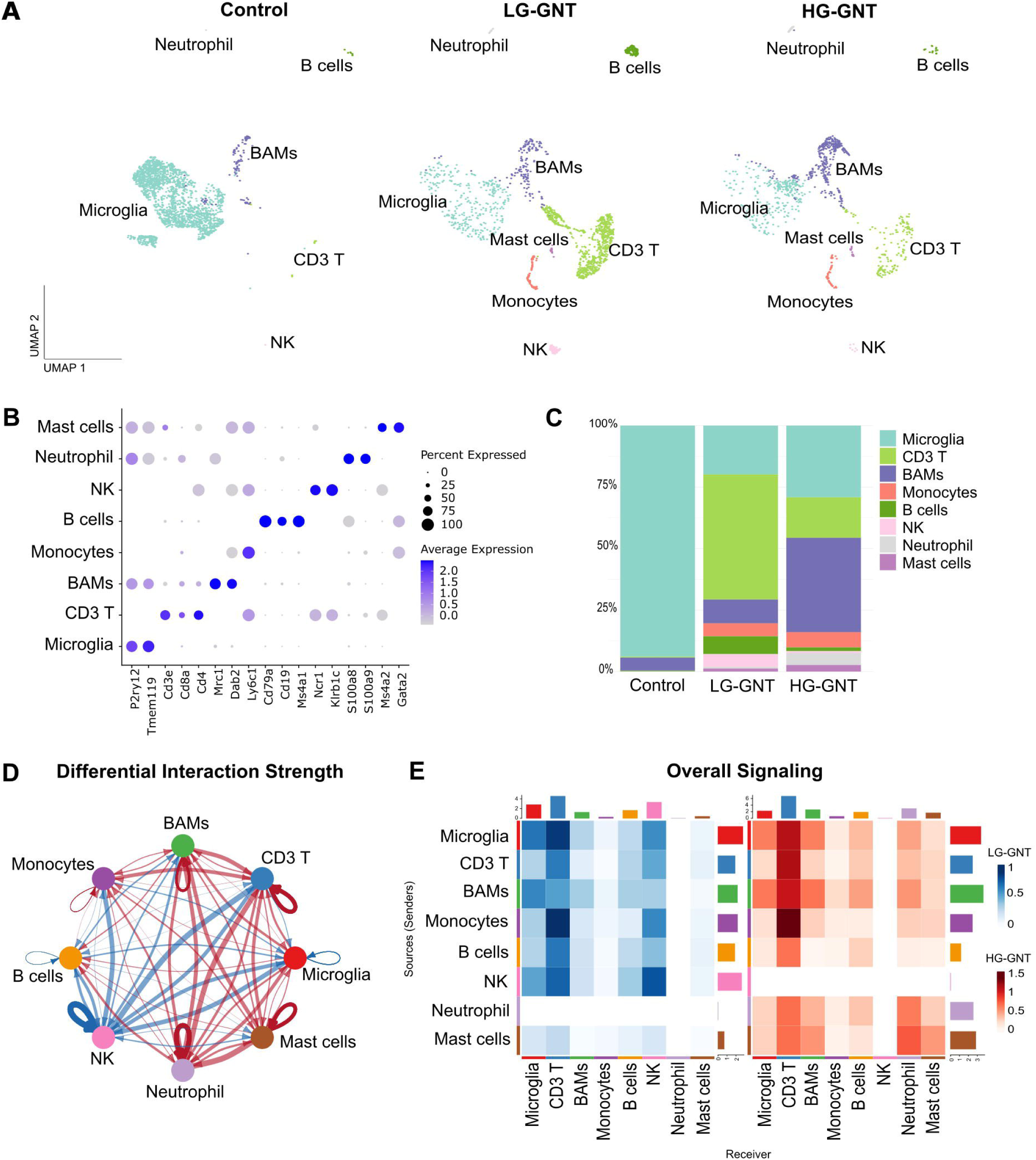
Single-cell resolution reveals a grade-dependent shift from lymphoid to myeloid dominance and intercellular communication in the GNTs microenvironment. **(A)** UMAP embeddings of the sub-clustered immune cell compartment, split by condition (Ctrl, LG-GNT, and HG-GNT). The analysis identifies distinct innate and adaptive immune populations, including microglia, BAMs, monocytes, CD3^+^ T cells, B cells, NK cells, neutrophils, and mast cells. **(B)** Dot plot showing the expression levels of canonical marker genes used to annotate the immune cell subpopulations. Dot size indicates the percentage of cells expressing a given gene, and the color intensity reflects the average expression level. **(C)** Stacked bar plot showing the relative frequencies of immune subpopulations within the total immune compartment across Ctrl, LG-GNT, and HG-GNT conditions. **(D)** Circle plot illustrating the differential interaction strength of inferred cell-cell communication networks among the immune populations as inferred by CellChat. This differential network highlights alterations between the LG-GNT and HG-GNT microenvironments, where red edges represent signaling interactions that are enriched in HG-GNT, and blue edges indicate signaling interactions enriched in LG-GNT. Edge thickness reflects the magnitude of differential signaling strength. The interaction weights, representing the sum of communication probabilities, were compared between LG-GNT and HG-GNT using CellChat’s differential analysis pipeline (p < 0.05). **(E)** Heatmaps showing overall signaling patterns (sum of interaction strengths) among immune cell populations in LG-GNT (left, blue scale) and HG-GNT (right, red scale). Rows correspond to sender (source) cell types, and the columns to receiver (target) cell types. The top and right bar plots represent the total incoming and outgoing signaling strength for each cell type, respectively.

Comparative analysis of immune cell type proportions revealed marked grade-dependent differences in immune composition across Ctrl, LG-GNT, and HG-GNT. Quantitative assessment confirmed that the immune population in the Ctrl group was composed of homeostatic microglia (93.4%), representing a resting, non-activated state of the CNS under physiological conditions, with a small presence of BAMs (5.3%). In contrast, the immune landscape of LG-GNT was predominantly lymphocyte-rich, with CD3^+^ T cells comprising the largest population (51%), highlighting a robust adaptive immune response (**Fig. 2C**). Conversely, the HG-GNT exhibited a markedly myeloid-dominant architecture, characterized by the predominance of innate populations such as BAMs and microglia, which accounted for 38.35% and 29.20% of immune cells, respectively (**Fig. 2C**). Moreover, while the fraction of CD3^+^ T cells, NK cells and B cells was lower in HG-GNT compared to LG-GNT, the neutrophils, largely absent in LG-GNT, were present within the HG-GNT TME (4.8%, **Fig. 2C**), indicating a distinct inflammatory landscape.

To investigate the intercellular signaling networks shaping the immune microenvironment, we performed CellChat analysis, which infers intercellular communication networks based on the gene expression of known ligand-receptor pairs between distinct cell populations. This analysis demonstrated intricate communication networks in both LG-GNT and HG-GNT (**Fig. 2D**), with multiple immune cell types actively engaged in intercellular crosstalk. Sender-receiver signaling analysis identified CD3⁺ T cells as the primary targets of intercellular communication in both LG-GNT and HG-GNT, predominantly receiving signals from infiltrating monocytes, microglia, and BAMs (**Fig. 2E**).

Taken together, these findings highlight a tumor grade-dependent remodeling of the immune TME characterized by a lymphocyte-rich profile in LG-GNT versus a complex myeloid-enriched environment in HG-GNT, yet both grades exhibit intricate intercellular signaling networks.

### T cell surveillance in low-grade GNT

Given the functional heterogeneity of T cells and their context-dependent roles in tumor immunity, we sought to characterize the T cell phenotypes within the TME (Pu et al., 2025). To this end, we sub-clustered the CD3^+^ T cell population. As highlighted above, the T cells were absent in the Ctrl group and only found in the GNTs, with a predominance in the LG-GNT. In both tumor entities, transcriptional profiling of T cells was composed of both CD8^+^ (*Cd8a*) and CD4^+^ (*Cd4*) T cells (**Fig. 3A-B**). Since we observed distinct cell abundance, we performed DEG analysis comparing the persisting T cells in HG-GNT versus LG-GNT. The results revealed minimal changes, with the T cells in HG-GNT showing higher upregulation of *Hspa1a* (HSP70), a major stress-inducible chaperone protein, and *Ccl4*, a pro-inflammatory chemokine associated with innate immune cell recruitment (**Fig. 3C**) [25].

**Figure 3:**
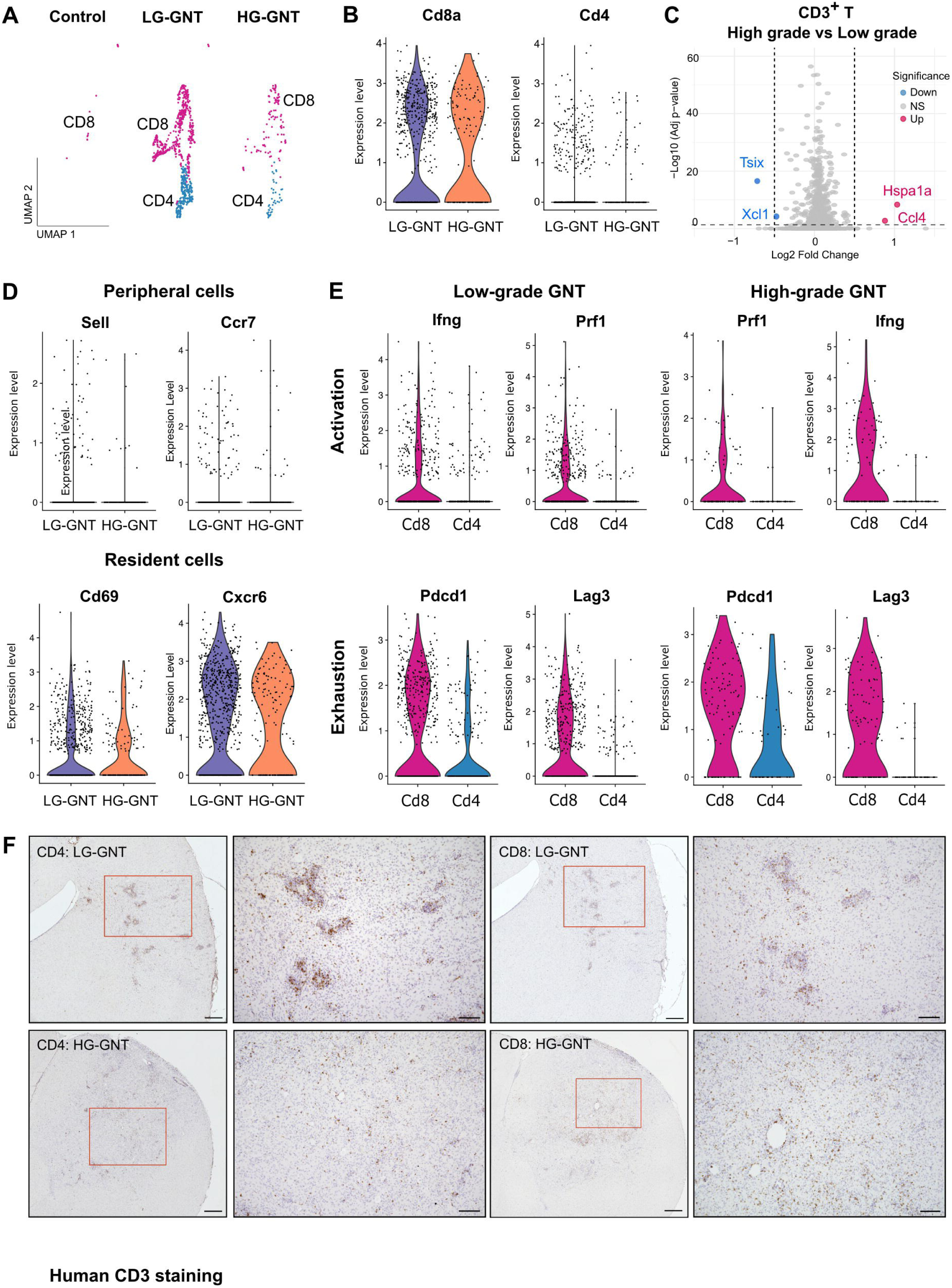
Transcriptional and histological characterization of T cells reveals tissue-resident memory features, chronic activation, and grade-dependent presence in GNTs. **(A)** UMAP embeddings of the CD3^+^ T cell compartment, split by condition (Ctrl, LG-GNT, and HG-GNT) and colored by annotated CD4^+^ and CD8^+^ T cell population. Of note, LG-GNT TME shows a higher abundance of CD4⁺ and CD8⁺ T cell subclusters compared to HG-GNT. **(B)** Violin plots depicting *Cd8a* and *Cd4* expression in T cells across LG-GNT and HG-GNT. **(C)** Volcano plot showing DEGs in CD3^+^ T cells between LG-GNT and HG-GNT. Significant DEGs are highlighted in red (up in HG-GNT) and blue (up in LG-GNT). **(D)** Violin plots detailing the expression of peripheral T cell markers (*Sell*, *Ccr7*, upper row) versus tissue-resident memory (Trm) markers (*Cd69*, *Cxcr6*, lower row). **(E)** Violin plots illustrating the expression profiles of activation and cytotoxic effector genes (*Ifng*, *Prf1*) as well as exhaustion markers (*Pdcd1*, *Lag3*) across CD8 and CD4 subclusters in LG-GNT (left) and HG-GNT (right), indicating chronic antigen stimulation. **(F)** Representative immunohistochemistry (IHC) images validating the distribution of CD4^+^ and CD8^+^ T cells in LG-GNT and HG-GNT sections. Zoomed-in insets (red squares) highlight the T cell infiltration in the LG-GNT and HG-GNT TME. (G) Representative IHC images demonstrating CD3^+^ T cell infiltration in human LG-GNT and HG-GNT tissue sections. Scale bars = 100 μm.

We next characterized the migratory and homing potential of the T cells. We found that T cells in the GNT TME, regardless of tumor grade, lacked expression of classical lymphoid homing markers such as *Sell* (gene encoding for CD62L) and *Ccr7* (**Fig. 3D**, top). Instead, they robustly expressed a signature characteristic of tissue-resident T cells (Trm), characterized by high expression of *Cd69* and the chemokine receptor *Cxcr6* (**Fig. 3D**, bottom). Although this Trm-like identity was shared between LG-GNT and HG-GNT, their functional state suggested chronic stimulation. In both tumor grades, T cells co-expressed cytotoxic effector genes, such as *Ifng* and *Prf1*, together with inhibitory checkpoint markers including *Pdcd1* (PD-1) and *Lag3* (**Fig. 3E**), indicating a state of chronic antigen stimulation.

To validate these single-cell findings, we performed immunohistochemical staining on tissue sections. Consistent with the scRNA-seq data, LG-GNT tissues exhibited a lymphocyte-rich phenotype, characterized by robust infiltration of both CD4^+^ and CD8^+^ T cells. Notably, these cells were not only restricted to the perivascular cuff but also found infiltrating the tumor parenchyma, suggesting an active surveillance mechanism. Conversely, T cells were detected at lower levels within HG-GNT tumors, which was in line with human LG-GNT and HG-GNT (**Fig. 3F-G**). Overall, these data define LG-GNT as an immunologically active tumor permissive to Trm infiltration, whereas HG-GNT exhibits restricted lymphocyte infiltration and diminished adaptive immune surveillance.

### Spp1-enriched immunosuppressive phenotype acquisition of microglia by transcriptional reprogramming in HG-GNT

Based on our GNT immune profile analysis (**Fig. 2A and 2C**), which revealed an expansion of the myeloid population within HG-GNT TME, we next sought to determine the grade-specific transcriptional heterogeneity of this key myeloid lineage. We first compared the myeloid profiles across Ctrl, LG-GNT and HG-GNT groups. Immunohistochemical staining for the microglia/ BAMs marker, IBA1, revealed strong morphological changes. While the cells in Ctrl tissues maintained a ramified, homeostatic morphology, those in both LG-GNT and HG-GNT exhibited an amoeboid phenotype characteristic of an activated state (**Fig. 4A**). Importantly, similar enrichment of infiltrating microglia was also detected in human GNT tissues, supporting the clinical relevance of our observations (**Fig. S2**). We further assessed the accumulation of CD206⁺ cells, a marker of macrophages. While absent in Ctrl tissue, CD206⁺ cells exhibited a pronounced infiltration in GNT tissue and were markedly enriched in the HG-GNT TME. (**Fig. 4B**).

**Figure 4:**
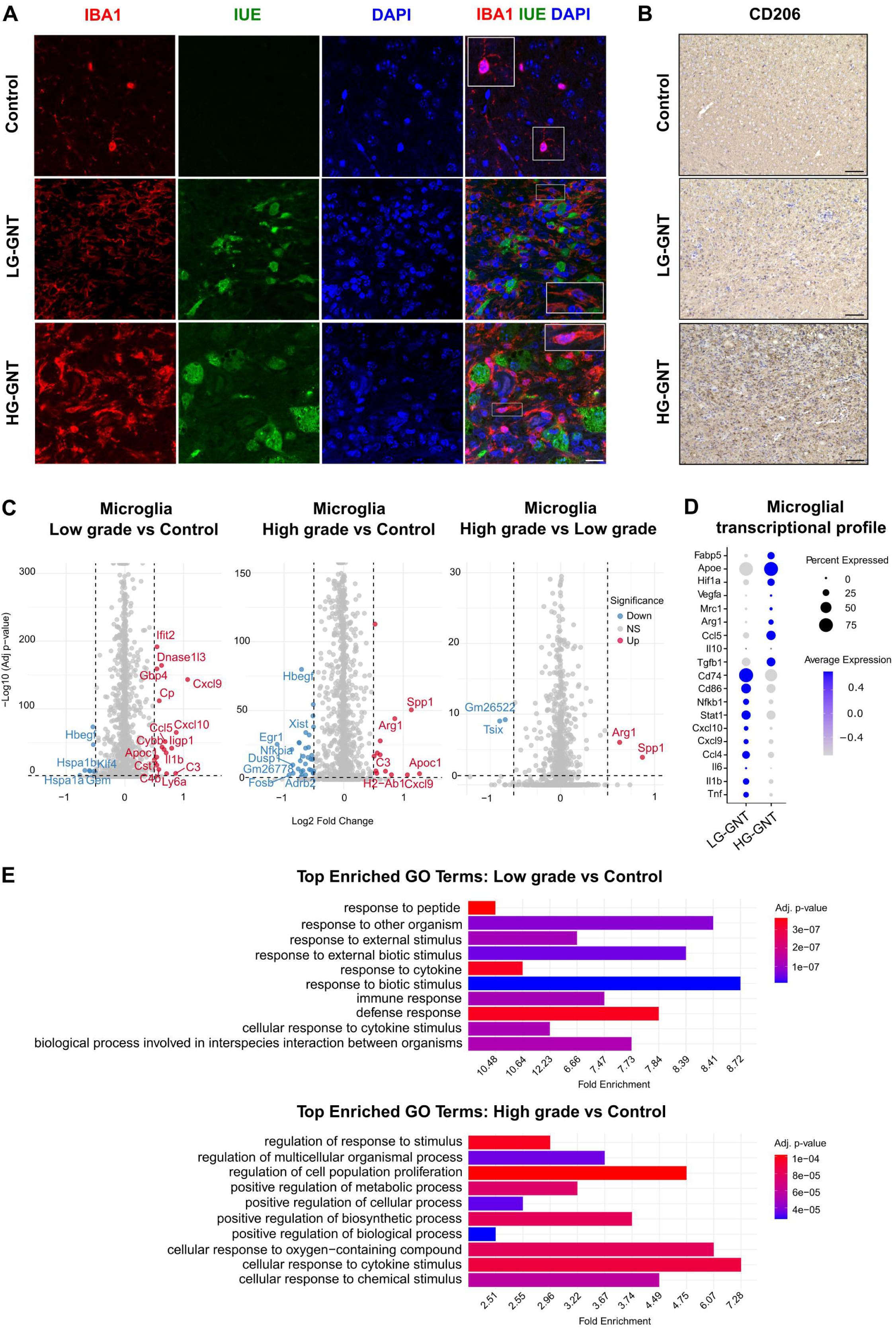
Microglia undergo a grade-dependent transcriptional rewiring from a pro-inflammatory to an immunosuppressive, tumor-supportive state. **(A)** Representative immunohistochemical (IHC) images from Ctrl, LG-GNT, and HG-GNT mouse tissue sections showing expression of IBA1-positive cells (red, marking microglia/BAMs), IUE-positive cells (green, identifying tumor cells), and DAPI (blue, nuclei). Insets (red boxes) display high-magnification views of IBA1-positive cells. Scale bars = 100 μm. **(B)** IHC staining for CD206 (*MRC1*), a marker of immunosuppressive macrophage activation, in Ctrl, LG-GNT, and HG-GNT sections. Note the marked enrichment of CD206^+^ cells specifically within the HG-GNT TME. Scale bars = 100 μm. **(C)** Volcano plots displaying DEGs in the microglial cluster. Comparisons are shown for LG-GNT vs Ctrl (left panel), HG-GNT vs Ctrl (middle panel), and HG-GNT vs LG-GNT (right panel). Red dots indicate significantly upregulated genes, while blue dots represent downregulated genes. Notably, LG-GNT microglia upregulate T cell-recruiting chemokines (e.g., *Cxcl9*, *Cxcl10*), whereas HG-GNT microglia are characterized by the robust upregulation of immunosuppressive markers such as *Spp1* and *Arg1*. **(D)** Dot plot showing expression of microglia-associated genes across LG-GNT and HG-GNT. Dot size indicates the percentage of cells expressing a given gene, and the color intensity reflects the average expression level. **(E)** Bar plots showing the top enriched Gene Ontology (GO) terms for biological processes in microglia from LG-GNT (top) and HG-GNT (bottom) compared to Ctrl. The x-axis represents the fold enrichment, and the color gradient indicates the statistical significance based on the adjusted p-value. Of note, LG-GNT microglia are enriched for terms related to active immune and defense responses, whereas HG-GNT microglia show enrichment for pathways driving cellular proliferation, metabolic regulation, and tissue remodeling.

To dissect the specific molecular alterations driving this myeloid-enriched TME, we then examined the transcriptomic changes within the microglia and BAMs population. DEG analysis of BAMs comparing LG-GNT with Ctrl revealed upregulation of activation and tissue-remodeling markers, such as *Mmp12*, *S100a4*, and *S100a6*. In contrast, comparison of HG-GNT with LG-GNT identified minimal transcriptional differences, with *Igkc* being the only notably upregulated gene. Given its limited functional relevance in this specific context, these findings suggest that the significance of BAMs in HG-GNT is driven by their marked accumulation rather than transcriptomic changes (**Fig. S3**). Together, these findings indicate that whereas BAMs expand substantially in number during malignant progression, they undergo relatively limited transcriptional remodeling. In contrast, microglia exhibit profound grade-dependent transcriptional reprogramming, identifying them as the principal immune population associated with the emergence of the high-grade state.

Interestingly, DEG analysis of microglia in both LG-GNT and HG-GNT versus Ctrl showed significant transcriptional differences. In the LG-GNT versus Ctrl comparison, augmented transcripts were primarily associated with immune defense and surveillance, including proinflammatory and interferon-stimulated genes (*Il1b, Ifit2*), and key chemokines involved in T cell trafficking (*Cxcl10, Cxcl9, Ccl5*). However, the HG-GNT versus Ctrl revealed a more complex landscape, characterized by the induction of genes involved in lipid metabolism (*Apoc1*) and immunosuppression (*Arg1, Spp1*), alongside persistent chemokine expression (*Cxcl9*). Furthermore, when directly comparing HG-GNT to LG-GNT microglia, DEG analysis identified *Spp1* (Osteopontin) as the most prominent and significantly upregulated gene in the HG-GNT TME. *Spp1* encodes a phosphoprotein known to dampen T cell responses and promote invasiveness in several tumor types (**Fig. 4C**) (Gu & Muller, 2025).

To understand the molecular context accompanying this robust *Spp1* augmentation more in depth, we analyzed microglial transcriptional profiles by assessing a targeted panel of genes related to immune responses, metabolism, and tissue remodeling, which collectively define microglial functional phenotypes. Our results showed that microglia in LG-GNT retained a proinflammatory signature, enriched for antigen presentation markers (e.g., *Cd74, Cd86, Nfkb1*) and chemokines associated with T cell recruitment, such as *Cxcl10, Cxcl9, and Ccl4*. Conversely, HG-GNT microglia exhibited a transition toward an immunosuppressive and tissue-remodeling phenotype. This state was characterized by the reduced expression of inflammatory markers and the distinct augmentation of *Apoe*, *Hif1a*, *Vegfa*, *Arg1*, and *Il10* mRNAs. Together, these transcriptional changes highlight a profound functional difference of microglia between the benign and malignant GNT variants (**Fig. 4D**).

Furthermore, this transcriptional divergence was reflected at the functional pathway level as revealed by gene ontology (GO) enrichment analysis. Microglia in LG-GNT were primarily enriched for immune and defense response pathways. In contrast, microglia in HG-GNT showed enrichment for pathways related to metabolic regulation, cell proliferation, and cellular response to chemical and cytokine stimulus (**Fig. 4E**). This transition further suggests that in the HG-GNT TME, microglia are functionally reprogrammed to support the metabolic and angiogenic demands of the tumor rather than being involved in anti-tumor defense.

Comprehensively, these data indicate a grade-dependent shift in the microglial compartment and identify *Spp1* as a putative key molecular driver of the high-grade microglial program, prompting further investigation as follows.

### SPP1 promotes microglia activation and stimulates growth dynamics of high-grade GNT components

To translate our transcriptomic findings into a mechanistic understanding of immune cell behavior within the TME, we next aimed to investigate whether *Spp1* may act as a functional driver of the HG-GNT microenvironment. We first mapped its expression across all major cell types identified in our single-cell dataset. The analysis revealed that *Spp1* expression was largely restricted to the immune cell compartment, with negligible expression observed in other cell types (**Fig. 5A**), confirming that *Spp1* is not a tumor-intrinsic marker but rather a product of the immune microenvironment in the context of GNT. Moreover, *Spp1* was robustly expressed by both microglia and BAMs, particularly within the high-grade environment (**Fig. 5B and Fig. S4**).

**Figure 5:**
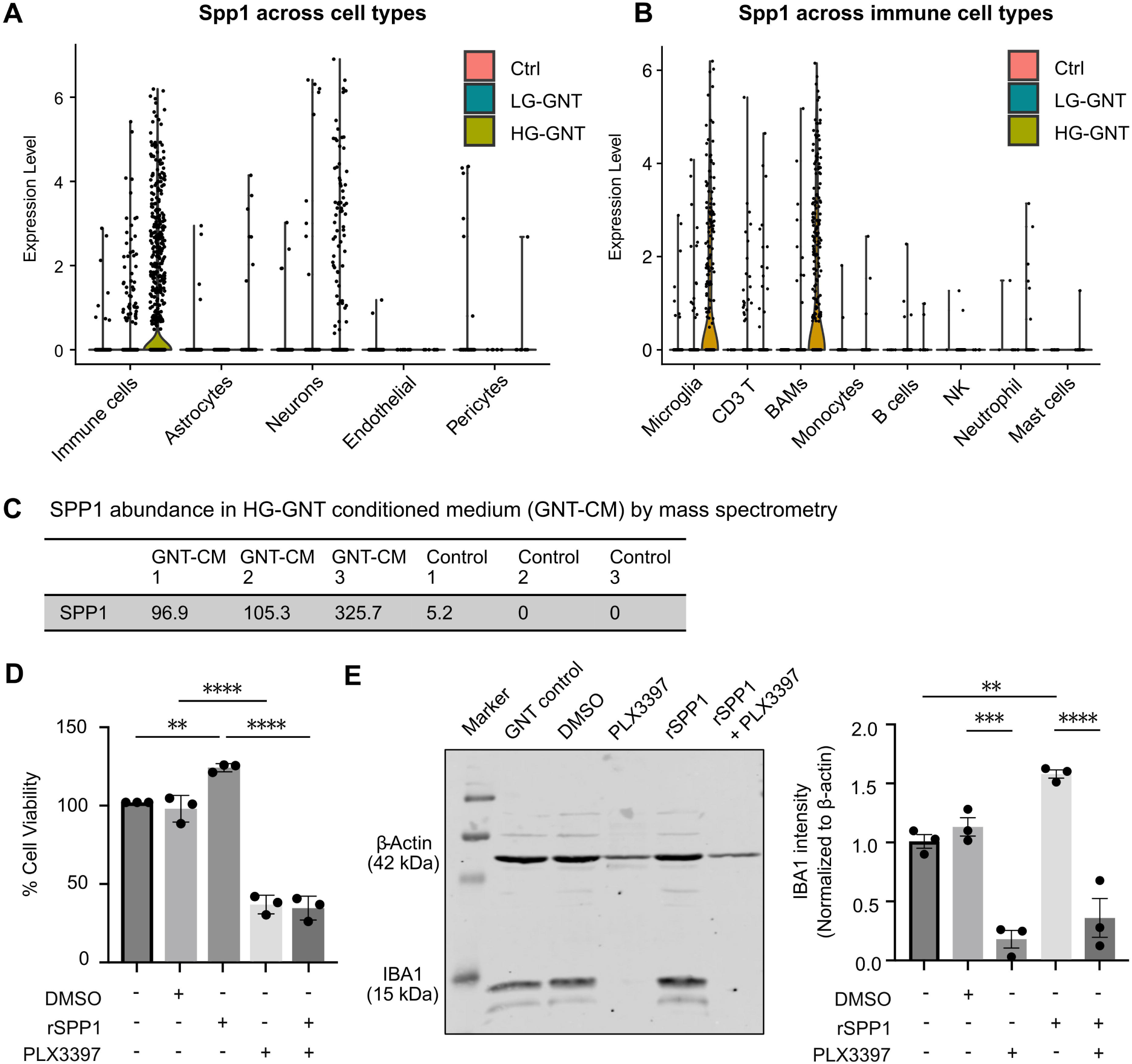
SPP1 is exclusively expressed by high-grade myeloid cells and drives tumor viability via a non-cell-autonomous, myeloid-dependent mechanism. **(A-B)** Violin plots displaying the expression levels of *Spp1* across (A) major cellular compartments and (B) distinct immune cell subpopulations across Ctrl, LG-GNT, and HG-GNT conditions. The data demonstrate that *Spp1* is uniquely restricted to the immune compartment and is predominantly expressed by microglia and BAMs specifically within the HG-GNT TME. **(C)** Table presenting the results from quantitative mass spectrometry analysis demonstrating the abundance of SPP1 protein in glioneuronal tumor-conditioned medium (GNT-CM) compared to control medium, confirming robust SPP1 secretion within the tumor milieu. (D) *Ex vivo* cell viability assay demonstrating the myeloid-dependent effect of SPP1 on tumor survival. Primary HG-GNT tumors were isolated, plated overnight, and subsequently treated for 72 hr. with vehicle (DMSO), PLX3397, recombinant SPP1 (rSPP1), or a combination of PLX3397 and rSPP1. The analysis reveals that while rSPP1 treatment significantly enhances overall cell viability, this pro-survival effect is completely abolished upon CSF1R-mediated myeloid depletion (PLX3397 + rSPP1). Data were normalized to the control within each biological replicate, and control viability was set to 100%. (** p < 0.01, **** p < 0.0001). **(E)** Representative Western blot (left) and corresponding protein intensity quantification (right) validating the modulation of the myeloid compartment. IBA1 protein levels were evaluated from the identically treated *ex vivo* cultures described in (D), with β-actin utilized as a loading control. The results confirm the highly efficient depletion of IBA1^+^ cells by PLX3397. Furthermore, the data illustrate that treatment with rSPP1 alone leads to a significant increase in the microglia/BAMs population. (** p < 0.01, *** p < 0.001, **** p < 0.0001).

We next aimed to verify whether this transcriptional upregulation translates to the secretion of functional protein. Mass spectrometry analysis of CM derived from primary HG-GNT cultures revealed a striking abundance of secreted SPP1 protein in HG-GNT-derived samples, whereas it was virtually undetectable in the controls (**Fig. 5C**). This implicitly assumes that GNT-associated microglia/ BAMs actively secrete SPP1 into the extracellular space.

To investigate the functional role of SPP1 in tumor growth, we used an *ex vivo* HG-GNT culture preparation as described before [26] and treated with recombinant SPP1 (rSPP1) and/or the CSF1R inhibitor PLX3397 to deplete microglia/BAMs. Cell viability assays revealed that rSPP1 significantly increased overall tumor culture viability compared to vehicle control. In contrast, treatment with PLX3397 markedly reduced viability, suggesting that the myeloid compartment is essential for GNT survival. Importantly, rSPP1 failed to rescue the viability loss induced by PLX3397, suggesting that its pro-tumoral effects depend on the presence of microglia/BAMs rather than acting directly on neoplastic cells (**Fig. 5D**). Accordingly, Western blot analysis showed increased IBA1 protein levels following rSPP1 treatment compared to control. Conversely, PLX3397 treatment completely abolished IBA1 protein levels, confirming effective depletion of microglia/BAMs (**Fig. 5E**). Together, these results demonstrate that SPP1 promotes tumor growth indirectly through microglia/BAMs-dependent mechanisms.

We next examined the receptor landscape of the tumor microenvironment to define the molecular signaling underlying these interactions (**Fig. S5**). Data from scRNA-seq analysis revealed robust expression of SPP1 receptors across multiple immune cell populations, including microglia and BAMs, indicating their capacity to directly receive and respond to SPP1 signaling within the HG-GNT TME. This intercellular crosstalk provides a mechanistic basis for the cell survival benefits observed in our functional assays. Taken together, these findings delineate a pathogenic axis in HG-GNT where myeloid-derived SPP1 acts as a key secreted factor that reinforces a tumor-supportive microenvironment. These data support a model in which malignant progression of GNTs is accompanied by a transition from T-cell-surveilled microenvironment toward SPP1-enriched myeloid state, driven by transcriptionally reprogrammed myeloid cells that promote tumor growth.

## Discussion

*BRAF*^V600E^-driven neuroectodermal neoplasms including primary brain tumors typically encounter significant immune cells [5, 7, 11, 34]. For *BRAF*^V600E^-driven GNTs, including GGs and PXAs, the molecular characteristics of immune cell infiltrates have remained vague so far.

Here, we provided new vistas of this pathogenetic aspect in a state-of-the-art GNT mouse model [6] that recapitulates key neuropathological features of human chronic epilepsy-associated neoplasms, particularly of GGs and PXAs (**Fig. S6**). In this model, immune activation is presumably orchestrated by the tumor-intrinsic hyperactivation of the PI3K/AKT/mTOR pathway, which has been linked to cellular metabolism and support of inflammatory cytokine production [28], thereby shaping a lymphocyte-permissive TME [41, 42].

This distinct *BRAF*^V600E^/mTOR-driven inflammatory signature appears to be fundamentally different from immune landscapes reported in diffuse glioma entities, which lack the strong association with chronic epilepsies. Particularly, IDH-wildtype gliomas exhibit massive immune infiltration that is predominantly biased toward immunosuppressive myeloid cells [1, 15]. In contrast, the present LG-GNT model appears to establish an immunogenic and permissive TME that supports prominent lymphoid infiltration. Therefore, the specific driver mutation in this LG-GNT fundamentally shapes the immune responses. While the KIAA1549-*BRAF* fusion in pilocytic astrocytomas relies on neoplastic CCL2 secretion to recruit tumor-supportive microglia [7], our *BRAF*^V600E^-driven model lacks prominent *Ccl2* upregulation. Instead, our single-cell data revealed that microglia act as the primary source of T cell attracting chemokines (e.g. *Cxcl9, Cxcl10, Ccl5*) and pro-inflammatory mediators (e.g. *Il1b, C3*) (**Fig. 4C**). This expression profile delineates a precise pathogenetic scenario for LG-GNT epileptogenesis. Although oncogenic *BRAF*^V600E^ intrinsically alters baseline neuronal excitability [17], it simultaneously triggers an epileptogenic neuro-glial crosstalk. While T cell infiltration is known to orchestrate phagocyte-mediated synaptic stripping via CCL2 [8], our data suggested an analogous but CCL2-independent mechanism whereby microglial CXCL9 and CXCL10 may recruit, retain and activate T cells [19], driving neuro-glial interactions that cause profound synaptic perturbation. Concurrently, microglial IL-1β and C3 promote neuronal hyperexcitability [40]. Thus, LG-GNT epileptogenesis emerges from a multifaceted mechanism, intrinsic *BRAF*-driven hyperexcitability compounded by an active immune response facilitating T cell-mediated synaptic loss and cytokine-induced epileptogenesis.

Importantly, epileptogenic mechanisms may be distinct in low-grade tumors and depend strongly on major aberrant pathway signaling in the individual neoplasms. In IDH1-mutant gliomas, the oncometabolite 2-hydroxyglutarate (2-HG) acts as an immune-independent epileptogenicity-promoting substrate by impairing glutamate biosynthesis and inducing mTOR-dependent hyperexcitability while simultaneously restricting immune infiltration [2, 23, 24] . This starkly contrasts with the highly immunogenic microenvironment of the present *BRAF*^V600E^-driven LG-GNT. Lacking a direct epileptogenic oncometabolite, network hyperexcitability in this tumor is fundamentally mediated by the outlined inflammatory cascades and MAPK/PI3K-induced immune activation.

In HG-GNT, the cellular and structural consequences of additional loss of *Trp53* apparently reshape the immune microenvironment. Beyond directly accelerating cell cycle progression, p53 loss induces rapid tumor growth that frequently outpaces blood supply, leading to necrosis, hypoxia, and possible disruption of the blood-brain barrier [3, 44]. Consequently, the release of damage-associated molecular patterns (DAMPs) from necrotic cells acts as a potent chemoattractant, promoting extensive immune cell recruitment [21]. Thus, the additional *Trp53* mutation establishes a fundamental dichotomy, facilitating the chaotic, DAMP-driven landscape dominated by immunosuppressive myeloid cells in the malignant GNT variant and thereby differentiating it from the localized, chemokine-driven lymphoid surveillance network in the benign GNT.

Importantly, our data suggest that malignant progression is not merely associated with quantitative changes in immune cell abundance but rather with a profound qualitative reorganization of the immune ecosystem. While LG-GNTs are characterized by a lymphocyte-rich microenvironment permissive to tissue resident T-cell surveillance, HG-GNTs exhibit extensive microglial reprogramming accompanied by expansion of the myeloid compartment. Notably, whereas BAMs predominantly increase in abundance, microglia undergo the most pronounced transcriptional remodeling, transitioning from a pro-inflammatory and T-cell-recruiting state toward an immunosuppressive phenotype enriched for *Spp1*, and *Arg1* expression. These findings identify microglial plasticity as a central feature of malignant progression in GNTs.

Importantly, our data suggest that malignant progression is not merely associated with quantitative changes in immune cell abundance but rather with a profound qualitative reorganization of the immune ecosystem. While LG-GNTs are characterized by a lymphocyte-rich microenvironment permissive to tissue resident T-cell surveillance, HG-GNTs exhibit extensive microglial reprogramming accompanied by expansion of the myeloid compartment. Notably, whereas BAMs predominantly increase in abundance, microglia undergo the most pronounced transcriptional remodeling, transitioning from a pro-inflammatory and T-cell-recruiting state toward an immunosuppressive phenotype enriched for Spp1, Apoe, Arg1, and Vegfa expression. These findings identify microglial plasticity as a central feature of malignant progression in GNTs.

Interestingly, while neoplastic cells are known to be a primary source of SPP1 in many other solid tumors [35, 43], we demonstrate that in HG-GNT, *Spp1* is predominantly expressed by microglia and BAMs, rather than by neoplastic cells. Our *ex vivo* experiments reveal that the addition of rSPP1 successfully enhanced overall tumor culture viability. However, upon CSF1R-mediated myeloid depletion, rSPP1 failed to increase this viability, indicating a strictly non-cell-autonomous mechanism. Despite the presence of cognate receptors in the TME that could theoretically enable direct signaling to tumor cells [12], SPP1 appears to primarily act on the myeloid compartment, promoting cell proliferation. These findings position the innate immune system not merely as a passive responder, but as an active driver of myeloid cell proliferation and activation [31]. This reinforcement is further supported by the distinct ontogeny of the BAMs within the HG-GNT TME compared to those in LG-GNT. We observed that the fraction of expanded BAMs population in HG-GNT exhibits a TIM4^−^ CCR2^+^ phenotype, but these markers were profoundly absent in the LG-GNT (**Fig. S7**). The absence of TIM4, a classical marker of yolk sac-derived tissue-resident macrophages, combined with high CCR2 expression, suggests a peripheral origin rather than a local expansion of resident cells [10]. Given its established role as a potent driver of monocytes/macrophages differentiation [27], we propose that an SPP1-rich milieu contributes to the polarization of infiltrating monocytes into tumor-supportive BAMs. This establishes a self-reinforcing feed-forward loop in which high-grade myeloid cells sustain their own activation and shape an immunosuppressive niche (**Fig. 6**). In conclusion, we define a distinct, grade-dependent immunological dichotomy in GNTs. We delineate the divergence between the T cell-enriched TME of low-grade tumors and an Spp1-associated myeloid niche of high-grade malignancies. The emerging pathogenetic scenarios in LG-versus HG-GNTs are as follows. In LG-GNTs, MAPK/PI3K hyperactivation is the determining factor promoting a microglia and T cell-enriched environment with strong epileptogenic effects, whereas in HG-GNTs, tumor progression is accompanied by a DAMP-associated and myeloid-enriched immune landscape. Collectively, our findings support a model in which malignant progression of GNTs is accompanied by a transition from a T-cell-surveilled microenvironment toward an SPP1-enriched myeloid state, in which transcriptionally reprogrammed microglia become dominant regulators of a tumor-supportive niche. Future studies will test treatment concepts targeting specifically the prepondering immune cell infiltrates in benign versus malignant GNTs.

**Figure 6:**
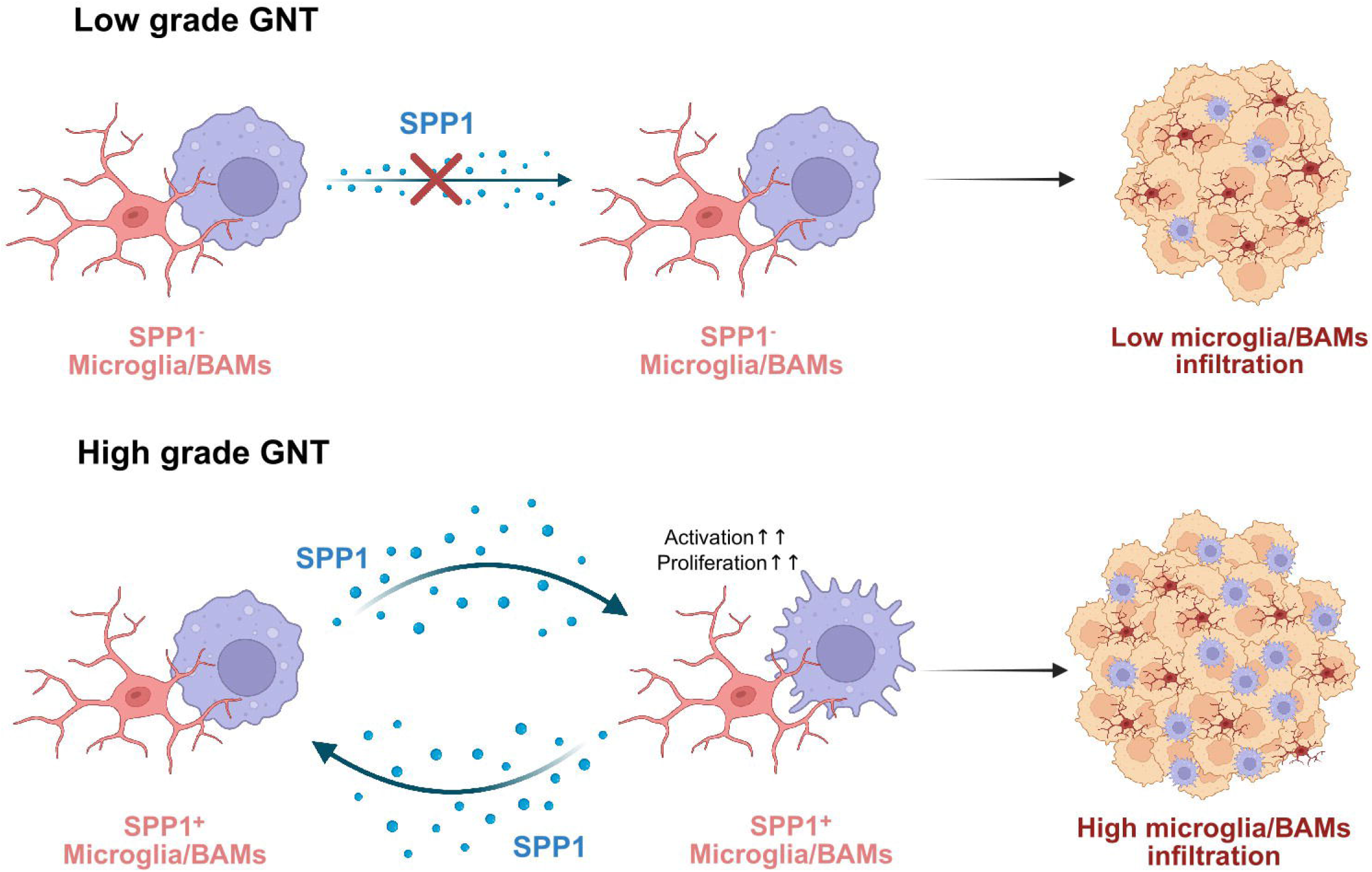
Schematic model illustrating the SPP1 - driven feed-forward loop mediating myeloid expansion in HG-GNTs. **(A)** LG-GNT; In the less aggressive state, the TME is characterized by SPP1-negative (SPP1^−^) microglia and BAMs. The absence of SPP1 signaling maintains a baseline level of myeloid infiltration, preserving a permissive niche that allows for active T cell surveillance. **(B)** HG-GNT; Upon progression to a high-grade state, the TME undergoes profound transcriptional rewiring. Microglia and BAMs robustly upregulate and secrete SPP1. This establishes a self-reinforcing, paracrine/autocrine feed-forward loop that intensely drives the activation and proliferation of the myeloid cells themselves. Ultimately, this mechanism enhances the massive expansion of the myeloid compartments, thereby shaping a hostile and lymphocyte-excluded microenvironment that physically and functionally restricts adaptive anti-tumor immunity.

## Acknowledgments

We thank Sabine Opitz, Pia Trebing and Shayne Gilgenbach for excellent technical assistance. This work was supported by the BONFOR program and the Epilepsy Surgery Biobank of the Medical Faculty at the University of Bonn. Protein identification was performed at the Core Facility Analytical Proteomics, University of Bonn, University Hospital Bonn, Institute of Biochemistry and Molecular Biology. Funded by the Deutsche Forschungsgemeinschaft (DFG, German Research Foundation) – Projektnummer 386936527. The work related to single cell transcriptomics was supported by the Deutsche Forschungsgemeinschaft (DFG) Research Infrastructure West German Genome Center (project ID 407493903) as part of the Next Generation Sequencing Competence Network (project 423957469). Next Generation Sequencing analyses were carried out at the production site Bonn (West German Genome Center Bonn/Next Generation Sequencing Core Facility of the Medical Faculty at the University of Bonn).

## Funding

Our work was supported by the Deutsche Forschungsgemeinschaft/German Research Council (to AB (BE 2078/10-2, project P7 in FOR 2715 ‘Epileptogenesis in Genetic Epilepsies’) and to SS (SCHO 820/6-2, SCHO 820/7-1, SCHO 820/8-1, SCHO 820/9-1, SCHO 820/10-1)) and by BONFOR (AB, SS, SC).

## Author contributions

**Kittitach Sri-ngern-ngam:** Conceptualization, Methodology, Validation, Analysis, Investigation, Data curation, Writing – original draft, Writing – review & editing, Visualization, Project administration.

**Philipp Müller:** Conceptualization, Methodology, Validation, Formal analysis, Investigation, Data curation, Visualization.

**Anne Quatraccioni:** Conceptualization, Methodology, Validation, Supervision.

**Valentina Zschernack:** Formal analysis, Investigation, Data curation, Visualization.

**Motaz Hamed:** Investigation, Resources.

**Rainer Surges:** Conceptualization, Supervision.

**Susanne Schoch:** Conceptualization, Funding acquisition.

**Julika Pitsch:** Conceptualization, Project administration.

**Albert J. Becker:** Conceptualization, Writing – original draft, Writing – review & editing, Supervision, Project administration, Funding acquisition.

**Silvia Cases-Cunillera:** Conceptualization, Methodology, Validation, Investigation, Writing – original draft, Writing – review & editing, Visualization, Supervision, Project administration.

## Data Availability

The data will be made available upon reasonable request.

## Declarations

### Conflict of interest statement

None of the authors has a conflict of interest.

### Ethical approval

All animal procedures were performed in accordance with European guidelines of the European Parliament and of the Council on the protection of animals used for scientific purposes, European Directive (2010/63/EU) and federal law (TierSchG, TierSchVersV) and approved by the country of North Rhine Westphalia (Landesamt für Natur, Umwelt und Verbraucherschutz, LANUV, Germany), and reported in compliance with the ARRIVE guidelines. All human biopsies were conducted in accordance with the Declaration of Helsinki. Informed written consent was obtained from every patient according to the approvals of the local ethics committee (Nr. 372/22).

## Notes

### Competing Interest Statement

The authors have declared no competing interest.

