## Supplementary_data for "Single-cell profiling reveals tumor grade-dependent immune remodeling in *BRAF*^V600E^-driven glioneuronal tumors"

Short title: Grade-dependent immune remodeling in glioneuronal tumors

*Corresponding author:
Silvia Cases-Cunillera
Department of Epileptology
University of Bonn Medical Center
Venusberg-Campus 1
53127, Bonn, Germany


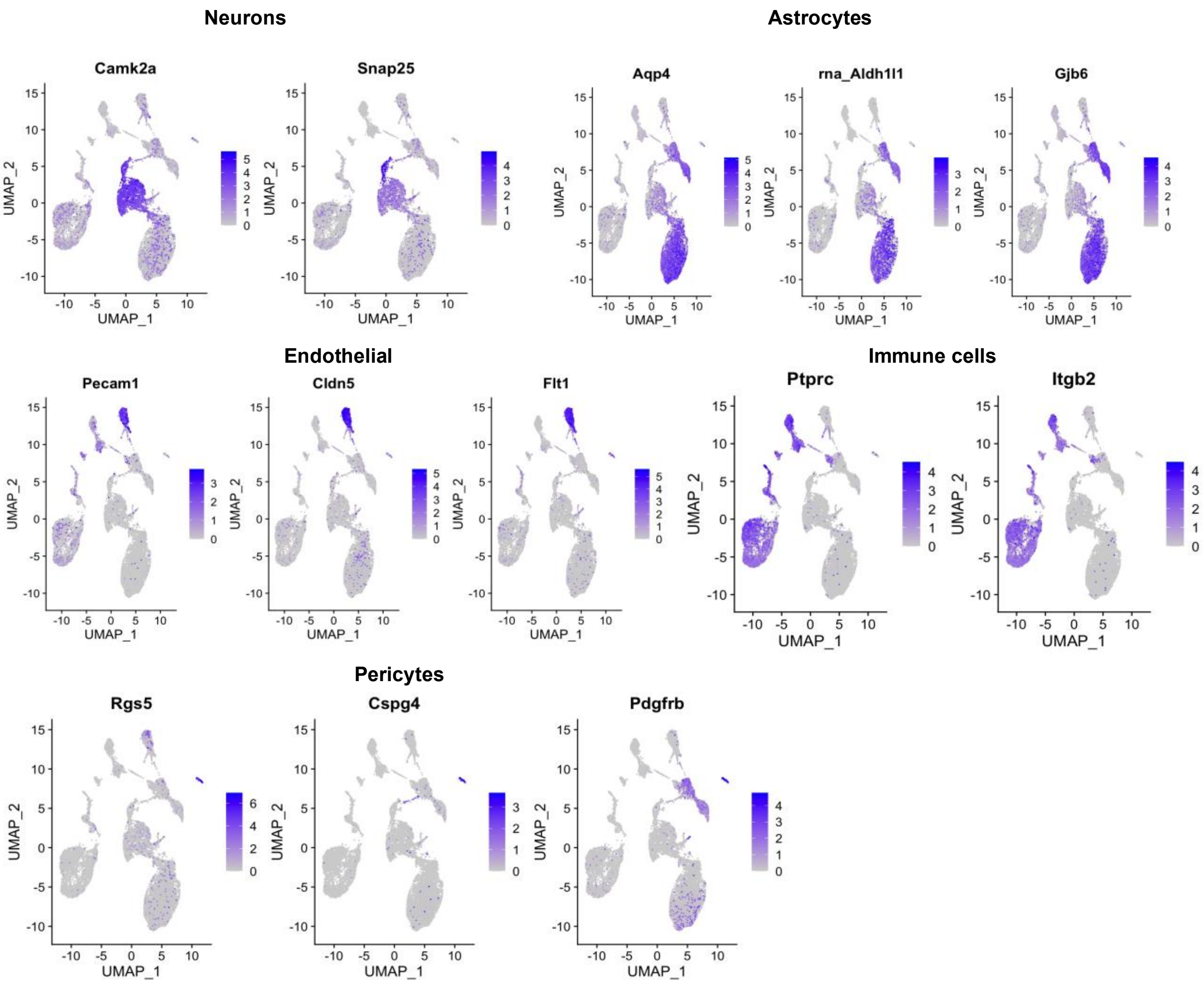


**Supplementary Figure S1. Expression of canonical marker genes across cell populations.** UMAP feature plots illustrating the normalized expression levels of established marker genes used for the annotation of distinct cell types. The panels display the expression of: Neurons: *Camk2a* and *Snap25*, Astrocytes: *Aqp4*, *Aldh1l1*, and *Gjb6*, Endothelial cells: *Pecam1, Cldn5*, and *Flt1*, Immune cells: *Ptprc* and *Itgb2* and Pericytes: *Rgs5, Cspg4*, and *Pdgfrb.*

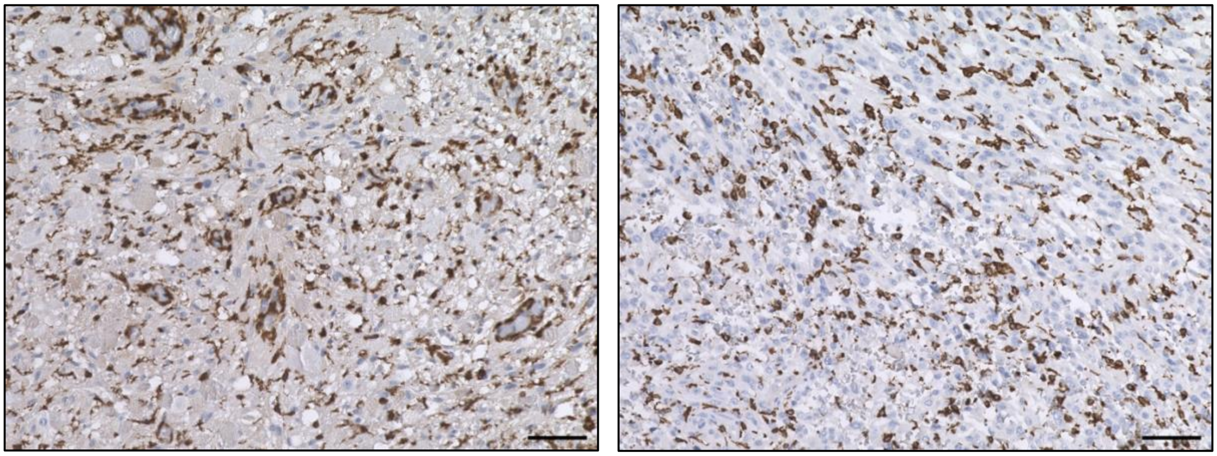

**Supplementary Figure S2: Microglial infiltration in human GNTs.** Representative images of IBA1 immunohistochemical staining highlighting microglia in human LG-GNT (left panel) and HG-GNT (right panel) tissues. Positive IBA1 staining is indicated by the brown color. Scale bar = 100 μm.

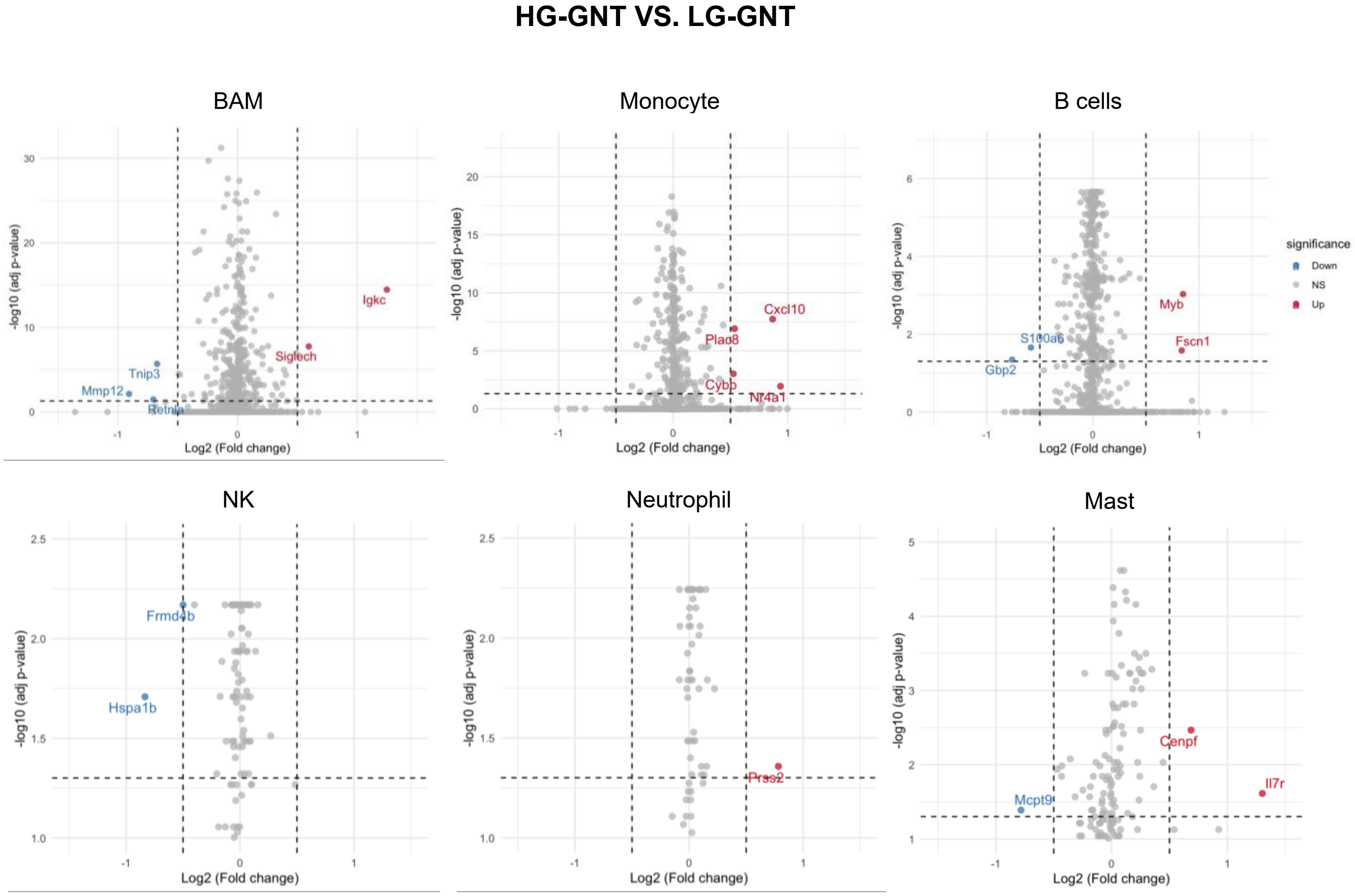

**Supplementary Figure S3: Differential gene expression in immune cell subpopulations between LG-GNT and HG-GNT**. Volcano plots displaying differentially expressed genes (DEGs) across distinct immune cell populations (BAMs: border-associated macrophages, Monocytes, B cells, NK: natural killer cells, Neutrophils, and Mast cells) comparing HG-GNT to LG-GNT. The x-axis represents the $\log_2$(fold change) and the y-axis represents the $-\log_{10}$ (adjusted p-value). Horizontal and vertical dashed lines indicate the statistical significance thresholds (adjusted $p < 0.05$ and $|\log_2 FC| > 0.5$, respectively). Significantly upregulated and downregulated genes in HG-GNT are highlighted in red and blue, respectively, while non-significant genes are shown in gray.

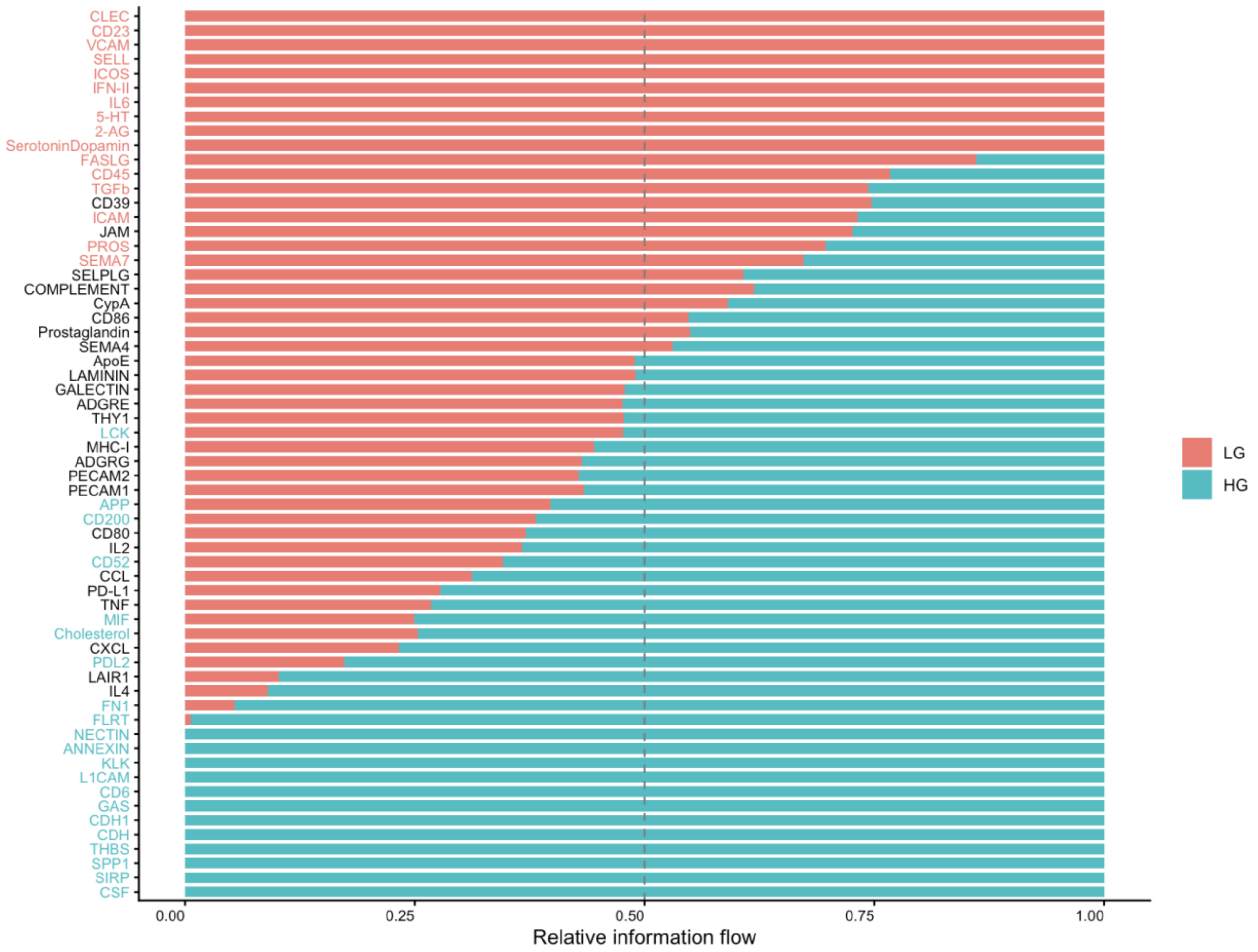


**Supplementary Figure S4: Comparison of overall signaling pathway information flow between LG-GNT and HG-GNT.** Stacked bar chart illustrating the relative information flow of inferred signaling pathways comparing LG-GNT (orange) and HG-GNT (blue), analyzed via CellChat. The overall information flow is defined as the sum of communication probabilities among all identified cell populations within the network.

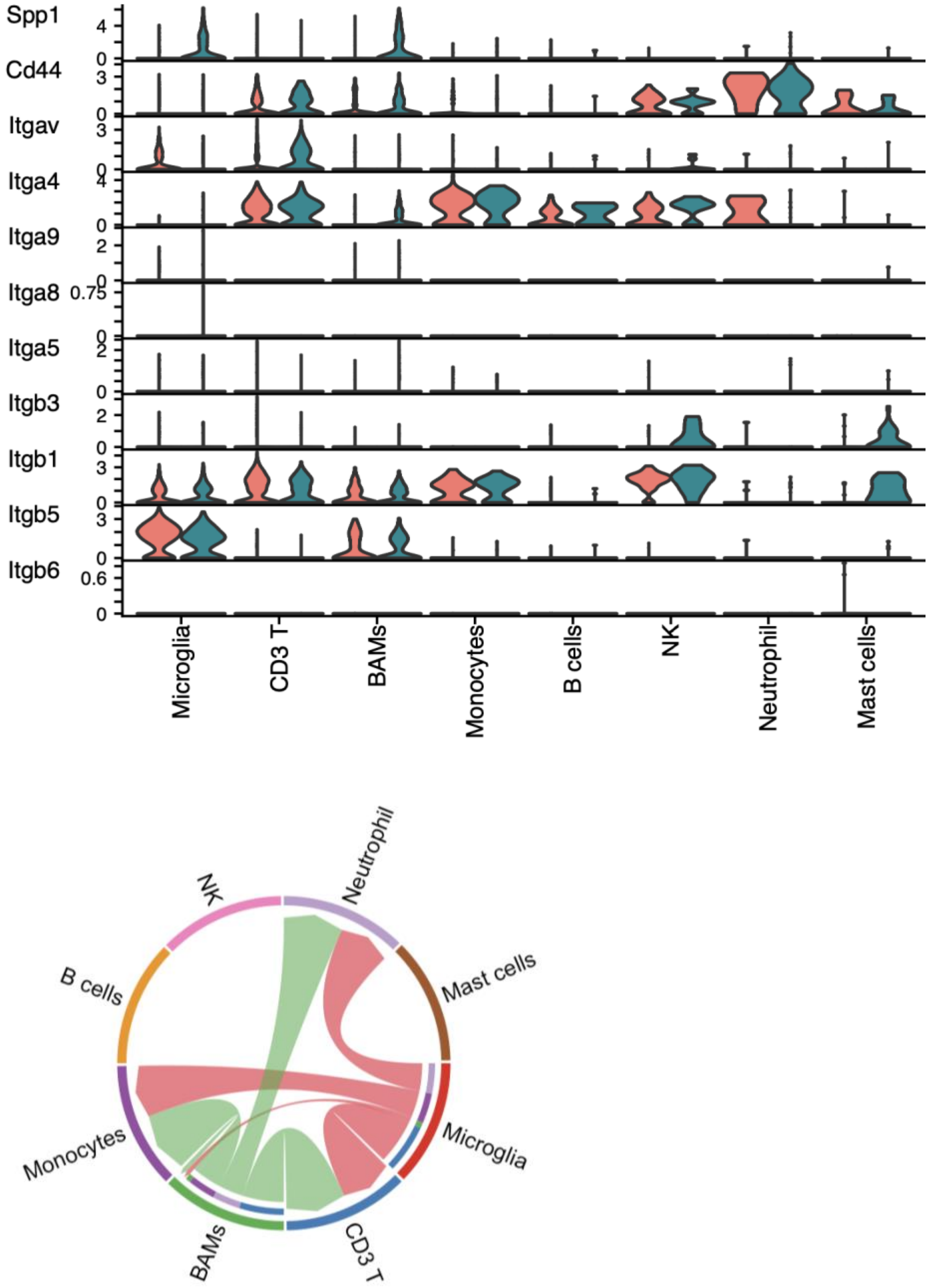


**Supplementary Figure S5: SPP1 signaling pathway communication network and expression of its ligand-receptor pairs. (Top)** Stacked violin plots displaying the expression distribution of the *Spp1* ligand and its cognate receptors (*Cd44* and various integrin subunits) across distinct immune cell populations. The colors indicate the comparison between the two groups (organge for LG-GNT and green for HG- GNT).

**(Bottom)** Chord diagram illustrating the inferred intercellular communication network of the SPP1 signaling pathway among the identified immune cell types. The width of the edges indicates the communication probability (strength) between the interacting cell populations.

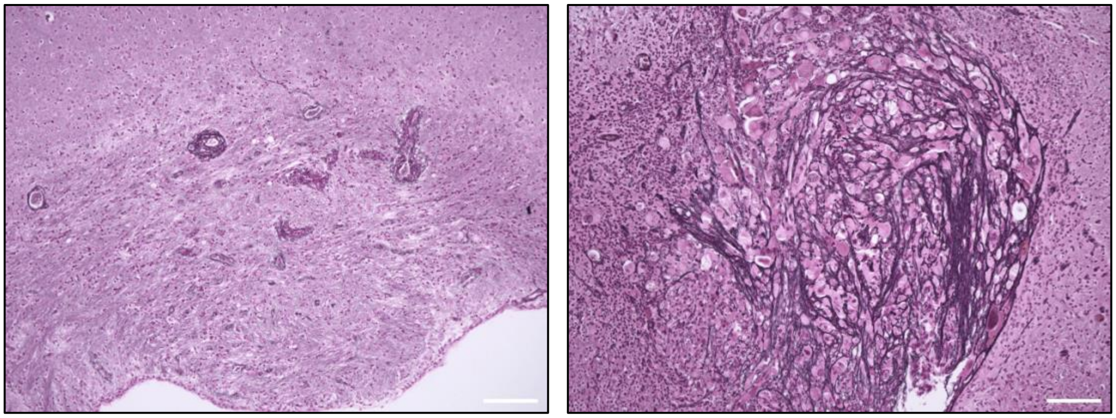

**Supplementary Figure S6: Reticulin fiber network in mouse GNTs.** Representative images of reticulin staining in tissue sections from mouse LG-GNT (left panel) and HG-GNT (right panel). The stain highlights the reticulin fiber network (type III collagen) as dark black/brown linear structures. A markedly increased density and complexity of the reticulin network is observed in the HG- GNT compared to the LG-GNT counterpart. Scale bar = 100 μm.

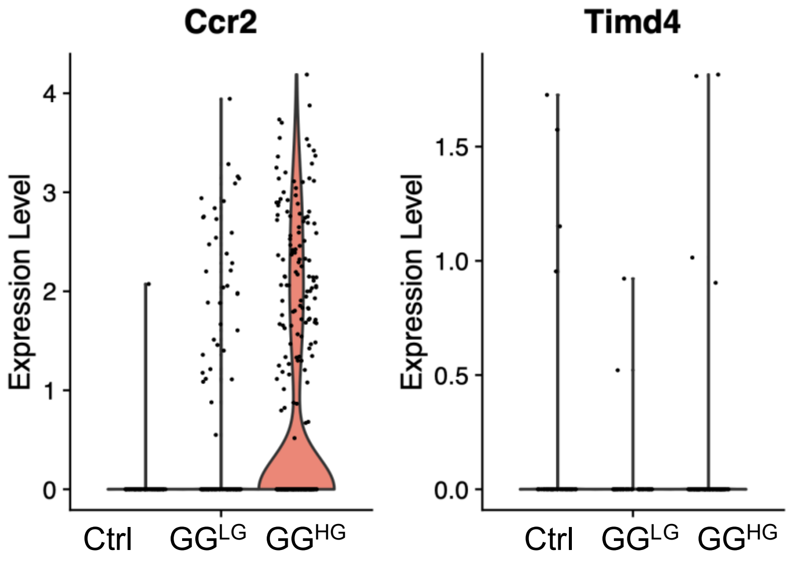


**Supplementary Figure S7: Expression profile of ontogeny markers indicating a monocyte-derived origin for BAMs.** Violin plots illustrating the expression levels of the monocyte-derived marker *Ccr2* and the tissue-resident marker *Timd4* within the BAMs population. The expression is compared across Ctrl, LG-GNT and HG-GNT. The predominant Ccr2$^{+}$/Timd4$^{-}$ phenotype observed in the HG-GNT suggests that these BAMs primarily originate from the infiltration of peripheral blood monocytes rather than being tissue-resident macrophages. Each dot represents an individual cell.
